# Brain Segregation and Integration Relate to Word-Finding Abilities in Older and Younger Adults

**DOI:** 10.1101/2024.07.11.603138

**Authors:** Elise J. Oosterhuis, Neil Bailey, Kate Slade, Patrick J. C. May, Helen E. Nuttall

## Abstract

Previous research has shown that word-finding difficulties in older age are associated with functional and structural brain changes. However, the use of functional brain networks, measured through electroencephalography, to predict word-finding in older and younger adults has not yet been investigated. This study utilised resting-state electroencephalography data (61 channels) from the Leipzig Study for Mind-Body-Emotion Interactions dataset (Babayan et al., 2019) to investigate the relationship between functional brain networks and word-finding ability in healthy younger and older adults. Graph theory-based measures in individualised delta, theta, alpha, and beta bands were computed to assess brain segregation and integration of 53 older (aged 59-77) and 53 younger right-handed adults (aged 20-35). Word-finding ability was quantified as the number of orally produced words during a semantic and letter fluency task. Multiple linear regression revealed that, in older adults, greater functional connectedness in the delta band was associated with lower semantic fluency. Irrespective of age, greater modularity in the alpha band was related to lower semantic fluency. A greater small-world index in the delta band was related to better semantic fluency, irrespective of age. Increased brain integration in the delta band corresponded to greater semantic fluency in older adults. Hence, word-finding ability seems to be related to brain segregation and integration specific to the frequency band, possibly indicating alterations in cognitive control or compensatory shifts to less functionally specific frequency bands. The article further provides a discussion on neural dedifferentiation, hyper-synchronisation, study limitations, and directions for future research.

## Introduction

In older age, people experience problems with lexical access, typically commencing around the age of 40 or 50 years (Kavé & Knafo-Noam, 2015). These problems manifest as word-finding difficulties (Burke & Mackay, 1997; Kavé & Knafo-Noam, 2015; Marini & Andreetta, 2016; Mortensen et al., 2006), and are one of the most prominent problems associated with cognitive ageing (Burke & Shafto, 2004). Previous studies have linked word-finding difficulties to both functional and structural brain changes in older adults (Meinzer et al., 2009; Stamatakis et al., 2011; Wierenga et al., 2008). Functional brain networks, both task-related and at rest, reflect the neurophysiological organisation of the brain (Bullmore & Sporns, 2009) and change with age due to deterioration of brain structure. Such changes could explain age-related decreases in cognitive performance, such as deterioration of memory (Bullmore & Sporns, 2009; Gaál et al., 2010; Zangrossi et al., 2021). However, the link between age-related changes in functional brain networks and the association with word-finding difficulties has not yet been investigated. The current study aimed to establish whether such a link exists in healthy older adults. We expect that such an investigation may inform future development of neurocognitive biomarkers associated with communicative ability.

The identification of functional brain networks via electroencephalography (EEG) offers a promising tool for investigating the effects of physiological ageing and identifying biomarkers for age-related pathology, including dementia (Valizadeh et al., 2019; Vecchio et al., 2020). Using graph theory, functional brain networks and measures of functional connectivity can be derived, such as the strength of synchronisation between neuronal populations (for methodology, see Bullmore & Sporns, 2009). Such measures indicate the brain’s efficiency or strength of information transfer between different brain regions (Bullmore & Sporns, 2009; Fries, 2005). Functional brain networks can be characterised in terms of segregation and integration, which underlie cognition (Sporns, 2013; Sporns et al., 2004). Functional segregation reflects neuronal communication between neighbouring brain regions, with more segregation reflecting a pattern where brain regions are more strongly connected with neighbouring nodes than more distant nodes, and less segregation reflecting the opposite pattern. Functional integration refers to the connections between modules, enabling the network to integrate information that is distributed over multiple brain regions (Sporns, 2013). A balance between segregation and integration leads to a small-world network, allowing for global efficacy of information transfer between brain regions (Achard & Bullmore, 2007; Bassett & Bullmore, 2006; Bullmore & Sporns, 2009; Watts & Strogatz, 1998). Studies using resting-state EEG show that the small-world index decreases with age (Gaál et al., 2010; Moezzi et al., 2019; Petti et al., 2016; Vecchio et al., 2014, 2020), due to decreases in functional segregation with age (Damoiseaux, 2017).

Age-related changes in functional brain networks have also been linked to decreases in cognitive performance, for example, in executive functioning and memory (Andrews-Hanna et al., 2007; Fleck et al., 2016). Moreover, higher segregation in older adults might relate to better memory ability (Chan et al., 2014; Zangrossi et al., 2021). Andrews-Hanna et al. (2007) argued that ageing is accompanied by the disruption of functional networks underlying higher-order cognitive functions. Since age-related declines in word-finding abilities have been previously linked to changes in brain structure and function (Meinzer et al., 2009; Stamatakis et al., 2011; Wierenga et al., 2008), it is possible that changes in functional brain networks also relate to word-finding difficulties in older age.

A relationship between age-related changes in functional brain networks and word-finding difficulties could be explained by the neural dedifferentiation hypothesis, which posits that brain regions and networks become less functionally specific to cognitive processes with age (Li et al., 2001). Moreover, age-related decreases in neurotransmitters, such as dopamine, reduce the efficiency of information transfer between brain regions (Koen & Rugg, 2019; Li & Lindenberger, 1999; Li & Rieckmann, 2014). This then causes neural dedifferentiation and consequently increases interindividual differences in cognitive performance (De Felice & Holland, 2018; Hultsch et al., 2002; Koen & Rugg, 2019). It is therefore proposed that neural efficiency and, hence, cognitive processes are optimal in younger adults (McIntosh et al., 2014). Goh (2011) hypothesised that both the differences in behaviour between younger and older adults and the age-related neural dedifferentiation are directly related to age-related changes in functional connectivity. Hence, age-related changes in functional brain networks seem to be associated with changes in cognitive performance.

Finally, age-related changes in functional brain networks have been found to be specific to EEG frequency band (e.g., Gaál et al., 2010; Micheloyannis et al., 2007; Smit et al., 2012; Vecchio et al., 2014) and different cognitive functions have been linked to certain EEG frequency bands (for an overview, see Başar et al., 2001). Delta activity is involved in inhibiting irrelevant responses and is important for internal concentration (Harmony, 2013; Mousavi et al., 2020). Greater delta power in younger adults and greater connectivity in the delta band in older adults have been linked to higher semantic fluency performance (Fleck et al., 2016; Mousavi et al., 2020). Hence, functional connectivity in the delta band might be important in supporting verbal fluency performance in older adults. Moreover, theta oscillations may play an important role in cognitive control, semantic-related processing, working memory, behavioural monitoring, and letter fluency (Cavanagh & Frank, 2014; Mousavi et al., 2020; Wang, 2010). Alpha band activity has been proposed to reflect attention, working memory, and inhibition (Başar et al., 1999; Jensen & Mazaheri, 2010; Klimesch et al., 2007; Stam, 2000). Lastly, beta band activity may play a role in working memory, decision-making, and lexical-semantic retrieval processes (Gola et al., 2013; Siegel et al., 2009; Weiss & Mueller, 2012).

To our knowledge, this is the first study investigating the relationship between functional brain networks and age-related changes in word-finding ability using EEG. We used data from the Leipzig Study for Mind-Body-Emotion Interactions (LEMON; Babayan et al., 2019) to investigate this relationship. First, we hypothesised that the decline in word-finding ability with age is linked to a decrease in the connectedness of functional brain networks. Specifically, we predicted a main effect of delta-band brain segregation on semantic fluency in older adults. Second, we hypothesised that the age-related decreases in brain segregation (which are reflective of neural dedifferentiation) are positively related to word-finding ability. That is, we predicted a positive main effect of segregation and small-world index on verbal fluency. Lastly, because individual variability in cognitive performance increases with age, with minimal variation between younger adults, and because of optimal neural efficiency in younger adults, we hypothesised that the relationship between word-finding ability and the connectedness of functional brain networks is absent in younger adults. Thus, we predicted that brain segregation does not predict verbal fluency in younger adults. The hypotheses, predictions, and experimental design were preregistered on the Open Science Framework website at https://osf.io/u6p42.

## Methods

### Participants

Data were obtained from the LEMON dataset (Babayan et al., 2019), which contains resting-state EEG recordings and psychological assessments of 153 younger adults, aged 20-35 years, and 74 older adults, aged 59-77 years (mean age and *SDs* are unavailable in the LEMON database to protect participants’ anonymity). All participants in the dataset were German speakers. We selected data for the older adult group based on the following criteria: the availability of EEG resting state and verbal fluency data; the participants being right-handed, not suffering from depression (i.e., Hamilton Depression Rating Scale score lower than 14), not having an alcohol or substance use disorder (i.e., Alcohol Use Disorder Identification Test, AUDIT, score < 8 and a negative result on the drug screening test). Based on these criteria, we included the data of 53 older adults (25 females) in the current study. Data from the younger adults were filtered based on the same criteria and, subsequently, a subset of 53 younger adults (21 females) were randomly selected to match the sample size of the older adult group (see Table 1). The sample size was based on an a priori power analysis using data simulation (Brysbaert & Stevens, 2018; DeBruine & Barr, 2021). More detailed information on the a priori power analysis can be found in the Supplementary Materials. Further information on the LEMON dataset can be found in Babayan et al. (2019).

**Table 1.** Sample Characteristics and Comparisons between Age Groups.

|  | Younger adults<br>(*N* = 53) | Older adults<br>(*N* = 53) | *t*-test |
| --- | --- | --- | --- |
| Sex | 21 females; 32 males | 25 females; 28 males | $t(104) = -0.78$,<br>$p = .438$ |
| Hamilton Depression Rating Scale | $M = 2.70$ ($SD = 2.44$) | $M = 2.28$ ($SD = 2.67$) | $t(104) = -0.80$,<br>$p = .427$ |
| AUDIT | $M = 3.30$ ($SD = 1.73$) | $M = 2.75$ ($SD = 1.79$) | $t(104) = -1.60$,<br>$p = .112$ |

### Materials

To quantify word-finding ability, we used the scores of the verbal fluency tasks (i.e., the Regensburger Wortflüssigkeitstest), which were available in the dataset. Within two minutes, participants had to generate as many German words as possible starting with the letter “s” (i.e. letter fluency) or words belonging to the category “animals” (i.e., semantic fluency). For the current study, we used the number of correctly produced words of both tasks that were generated within the first minute, which is the time limit commonly used in standard versions of the task (Shao et al., 2014).

### EEG Recordings and Pre-processing

Resting-state EEG was recorded for 16 minutes, with alternating 60-second blocks of eyes-closed and eyes-open conditions. Only the eyes-closed condition was analysed in the current study. The set-up consisted of 61 channels arranged according to the 10-10 international system, with one additional electrode recording the vertical electrooculogram to monitor eye movements (see Babayan et al., 2019, for more information on the EEG recording setup). Data were pre-processed using MATLAB R2018a (MathWorks) and Fieldtrip (Oostenveld et al., 2010). The pre-processing pipeline “Reduction of Electroencephalographic Artifacts” (RELAX), which makes use of both Fieldtrip and EEGLAB, a MATLAB toolbox (Delorme & Makeig, 2004), was used to clean the continuous EEG data with the RELAX_wICA_ICLabel setting since the effect of the default multi-channel Wiener Filter cleaning approach has not been tested for use prior to analysis of EEG connectivity (Bailey, Biabani, et al., 2023; Bailey, Hill, et al., 2023).

Before cleaning the data with RELAX, the raw EEG data were downsampled from 2500 Hz to 1000 Hz. A 1-45 Hz Butterworth bandpass filter was applied. The RELAX pipeline identifies noisy channels via the PREP pipeline algorithm (Bigdely-Shamlo et al., 2015). Data from these channels were subsequently removed. Data from the remaining noisy channels were further removed using the default settings from RELAX. The mean proportion of the EEG data removed due to noisy channels was 0.047 and data from 59 channels *(SD* = 3), on average, were left after this removal. The cleaned data were re-referenced to the robust average reference before running the Independent Component Analysis (ICA) with the FastICA algorithm (Hyvarinen, 1999) and reducing artifacts identified by ICLabel (Pion-Tonachini et al., 2019) using Wavelet Enhanced ICA (Castellanos & Makarov, 2006). After data cleaning, excluded channels were interpolated using spherical spline (Delorme & Makeig, 2004).

### Functional Brain Networks

After cleaning the data, we applied a lowpass filter of 30 Hz and segmented the continuous data into 12-second epochs with 50% overlap (for more information on optimal epoch length for the debiased weighted Phase Lag Index (dwPLI), see Fraschini et al., 2016; Miljevic et al., 2022). Segments of the data were visually inspected to check data quality. The average proportion of data removed due to bad epochs was 0.205 *(SD* = 0.193). The average number of remaining epochs was 98. Next, we obtained the cross-spectral densities of the alpha, beta, theta, and delta bands using a Fourier transformation using the multitaper method based on Hanning tapers (see Figure 1A).

**Figure 1.**
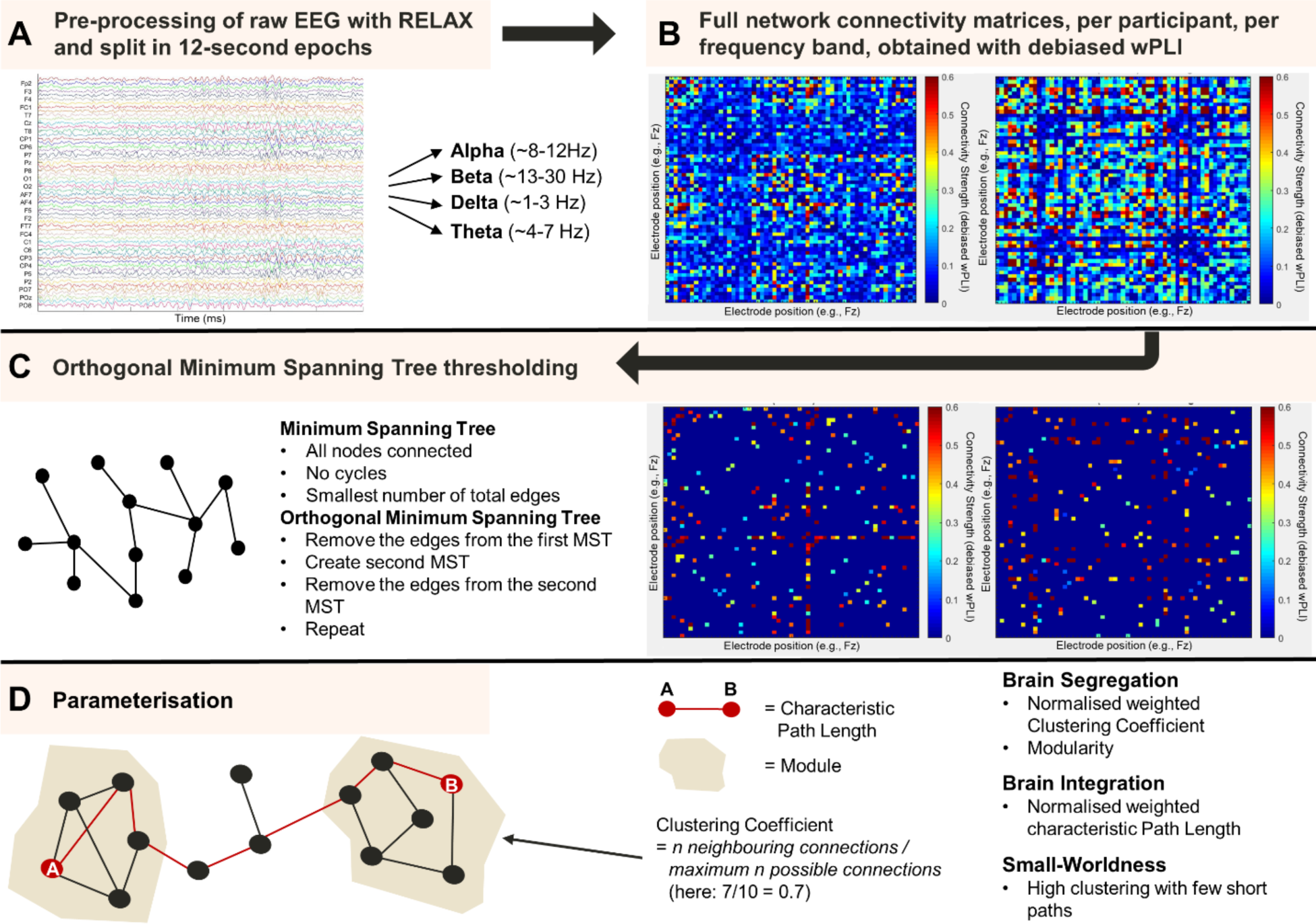
Processing Pipeline of the Resting-State Eyes-Closed EEG Data.

Studies have consistently shown that neurophysiological rhythms slow with ageing and that including conventional frequency bands can introduce a bias against older adults, also when conducting connectivity analyses (Chiang et al., 2011; Scally et al., 2018). Therefore, the alpha, beta, theta, and delta spectral boundaries were determined using the individual alpha peak frequency (IAPF). IAPF was calculated with eyes-closed resting-state EEG data using the restingIAF toolbox in Matlab (see Corcoran et al., 2018, for more information on how the IAPF was computed). The mean IAPF of older adults was 9.4 Hz *(SD* = 0.9 Hz) and 10.1 Hz *(SD* = 0.9 Hz) for younger adults. The alpha band was determined as IAPF -4 Hz to IAPF +2 Hz, theta as IAPF -6 Hz to IAPF -4 Hz, and delta as IAPF -8 Hz to IAPF -6 Hz (Babiloni et al., 2020; Klimesch, 1999). The beta band was determined as IAPF +2 Hz up to and including 30 Hz (Babiloni et al., 2020; Henelius et al., 2011).

To detect brain networks, the dwPLI was used to calculate the functional connectivity between all 61 channels, for each frequency per participant. Please, refer to the Supplementary Materials for the details on how dwPLI was computed. For the subsequent graph statistical processing steps and graph theoretical indices, we obtained the absolute values of the dwPLI to get an indication of the strength of connectivity between pairs of electrodes. The network construction resulted in a 61-by-61 weighted matrix for each frequency band and per participant (see Figure 1B).

We applied a data-driven method, namely the Orthogonalized Minimum Spanning Tree (OMST) algorithm to threshold the connectivity matrices (Dimitriadis et al., 2017). A minimum spanning tree is a graph with a minimum number of total edges, without cycles (i.e., the graph does not contain any loops), and where all nodes are connected (see Figure 5C). The OMST algorithm computes the minimum spanning tree (MST) over multiple iterations. These iterations are necessary as using a single MST might result in a graph that is too sparse for computing robust connectivity measures. For a more detailed explanation as to how the network graphs were thresholded using the OMST algorithm, please refer to the Supplementary Materials.

### Graph Theoretical Network Analysis

After thresholding, we applied graph theoretical analysis of the brain networks using Fieldtrip (Oostenveld et al., 2010) and the Brain Connectivity Toolbox (Rubinov & Sporns, 2010) in MATLAB (see Figure 1D). In the current study, the EEG sensors represent the nodes and the dwPLI values represent the weighted edges of the graph. Because we used weighted matrices, graphs for each frequency band were first normalised before computing the measures of brain segregation and integration, resulting in normalised weighted measures. All thresholded graphs were normalised by rescaling all weight magnitudes ranging between 0 and 1 (Bullmore & Sporns, 2009).

To quantify brain segregation (i.e., the clustering of functional networks into separate communities/groups), we calculated the weighted variant of the clustering coefficient (Onnela et al., 2005; Rubinov & Sporns, 2010) as well as of modularity (Newman, 2006; Rubinov & Sporns, 2010). Please, refer to the Supplementary Materials for the mathematical equations and descriptions of all connectivity measures. The clustering coefficient and modularity offer alternative statistics of brain segregation. Higher values of either statistic reflect greater local efficiency of information transfer in the brain (Bullmore & Sporns, 2009). That is, brain regions or communities are more specialised and have stronger connections within themselves (i.e., intra-community connections), facilitating efficient communication within those localised communities.

The degree of brain integration is quantified through the weighted version of the characteristic path length, which is the shortest path length between two nodes averaged across all node pairs (Rubinov & Sporns, 2010; Watts & Strogatz, 1998). Lower characteristic path length indicates greater global efficiency of information transfer in the brain. The clustering coefficient and modularity (brain segregation) and characteristic path length (brain integration) values were calculated for each participant for each frequency band separately. To examine the balance between brain segregation and integration, we computed the small-world index of each graph (Humphries & Gurney, 2008). Small-world indices with values higher than 1 indicate that the network is a small world (see the Supplementary Materials).

### Statistical Analysis

After obtaining the functional connectivity measures described above, the final data pre-processing and analysis were conducted in R (R Core Team, 2020). Following the preregistration, verbal fluency scores +/- 3 *SD*s would be considered outliers, however, no outliers were detected in the verbal fluency data. Missing values for all connectivity measures for OMST-thresholded networks can be found in Table 2. Full networks did not have any missing values. Even though the missing values in the OMST-thresholded graphs reduced the number of data points, the data will still provide important insights into the relationship between word-finding ability and age-related changes in functional brain networks. After the initial model fit, leverage points were identified as 2(number of predictors + 1)/number of observations, and subsequently removed to obtain the model’s best-fit.

**Table 2.**
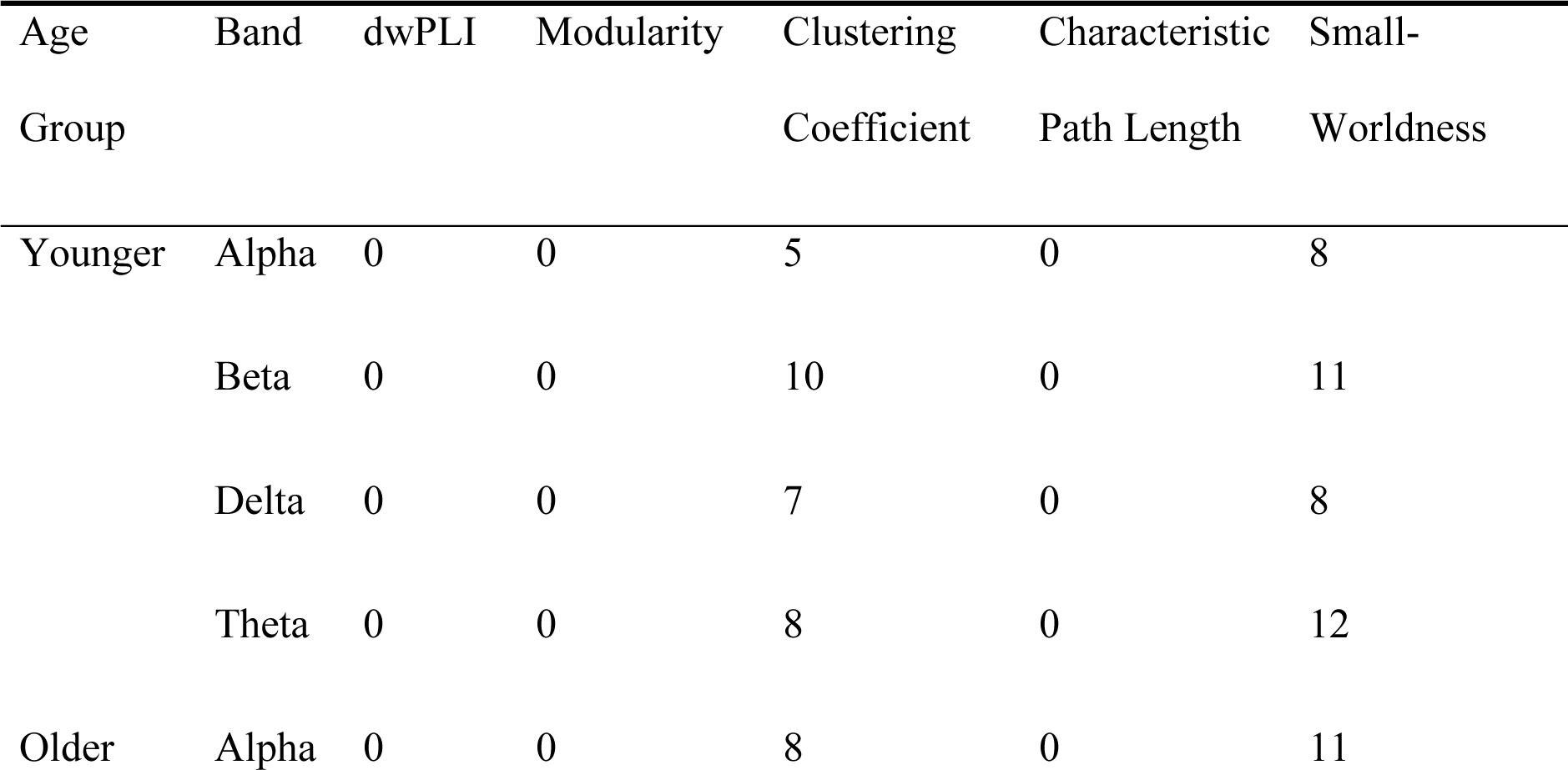
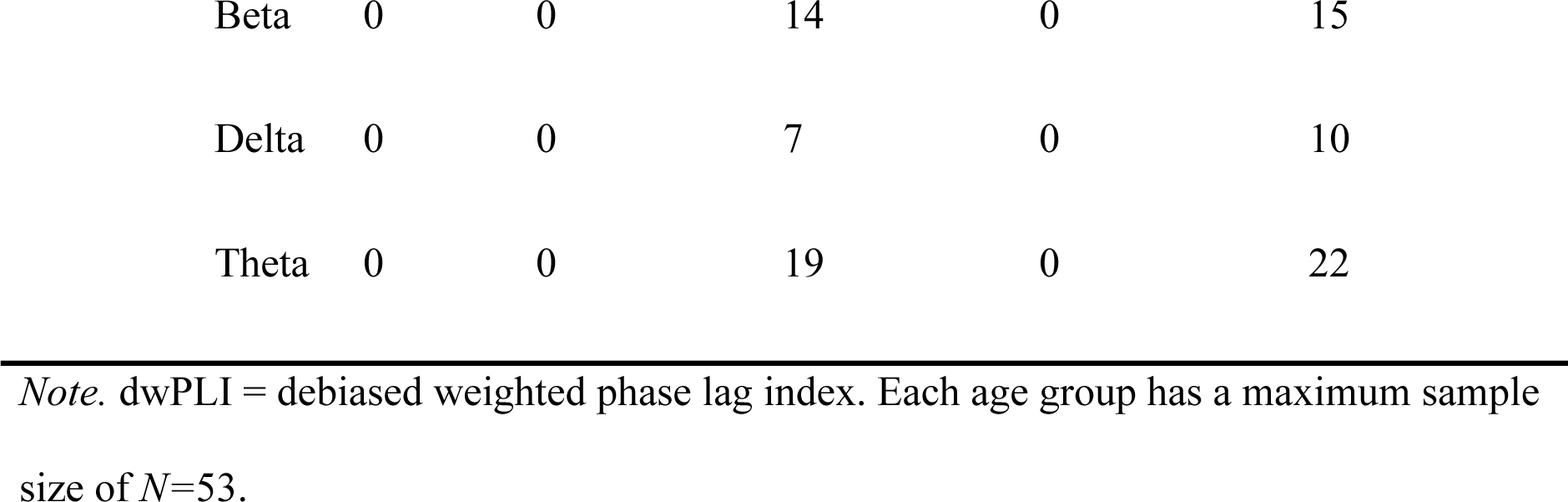
Missing Values for Each of the Connectivity Measures of the OMST-Thresholded Networks per Frequency Band.

To investigate whether age-related changes in brain segregation and integration are related to word-finding ability, we performed multiple linear regression analyses for each frequency band and each verbal fluency measure (i.e., letter and category fluency) separately. Because the networks were thresholded using a model-driven algorithm (i.e., OMST) and weighting was employed, we did not apply multiple corrections (which meets the criteria for optimal validity within a connectivity study, see the checklist by Miljevic et al., 2022). The outcome variables were the number of correctly produced words for the letter and category fluency tasks. To investigate the effect of age-related changes in brain segregation on verbal fluency, we included the interaction between clustering coefficient and age, and the interaction between modularity and age as predictors. To investigate the connectedness of the functional brain networks, we ran multiple linear regression analyses with the interaction between age and the small-world index as predictors of verbal fluency performance.

For brain integration, the predictor was the interaction between characteristic path length and age. All models included sex as a covariate because studies have shown that brain networks can differ between males and females (Foo et al., 2021). Both age and sex were contrast coded using “treatment contrasts” whereby younger adults were set as the reference level. All numerical predictors (i.e., the functional connectivity measures) were scaled for model interpretation. Assumptions of linearity, homoscedasticity, and normality of residuals were all met. Model diagnostics revealed the presence of leverage points for most models (i.e., values above 2(number predictors+ 1) / number of observations), which were subsequently removed (Cook, 1977). Only the results after the removal of leverage points are reported in the results section. This study was preregistered on the Open Science Framework (https://osf.io/u6p42). The quality of the connectivity analysis was checked against the checklist by Miljevic and colleagues (2022) and obtained a score of 5.5, which reflects high study quality.

### Deviations from the Preregistration

As preregistered, functional connectivity measures with values +/- 3 *SD*s from the mean were originally also considered as outliers. However, due to the sparsity of some functional brain networks, there were missing values in both age groups for the clustering coefficient and small-world index. To avoid reducing the dataset even further by removing outliers for the functional connectivity measures, we analysed the data with the detected outliers. Regarding brain segregation measures, the preregistration only mentioned the clustering coefficient as a brain segregation measure for hypotheses 1 and 2. However, since brain segregation is reflected by both clustering coefficient and modularity (Rubinov & Sporns, 2010), we included both the interaction between clustering coefficient and age, as well as the interaction between age and modularity in the statistical models. Such models would represent brain segregation better than solely including the clustering coefficient.

## Results

### Behavioural Data

Before analysing the functional brain networks, a behavioural difference in verbal fluency performance between younger and older adults was confirmed. For semantic fluency, older adults obtained a mean score of 22 correctly produced words *(SD* = 5.5; range = 12-37 words) and younger adults obtained a mean score of 25 correctly produced words *(SD*=5.1; range=15-39 words), and this difference between age groups was significant (*t(* 418.68=-5.80, *p* < .001). For letter fluency, older adults obtained a mean score of 13 correctly produced words *(SD*=3.1; range=5-19 words) and younger adults obtained a mean score of 15 correctly produced words *(SD*=3.5; range=7-23 words), and this difference between age groups was also significant (*t*(416.4)=-6.72, *p* < .001).

### Graph Analysis

To investigate the age-related changes in functional brain networks, the brain segregation and integration measures were computed for the alpha, beta, delta, and theta bands. Mean and standard deviations for the graph theoretical measures of the OMST-weighted graphs can be found in Table 3.

**Table 3.**
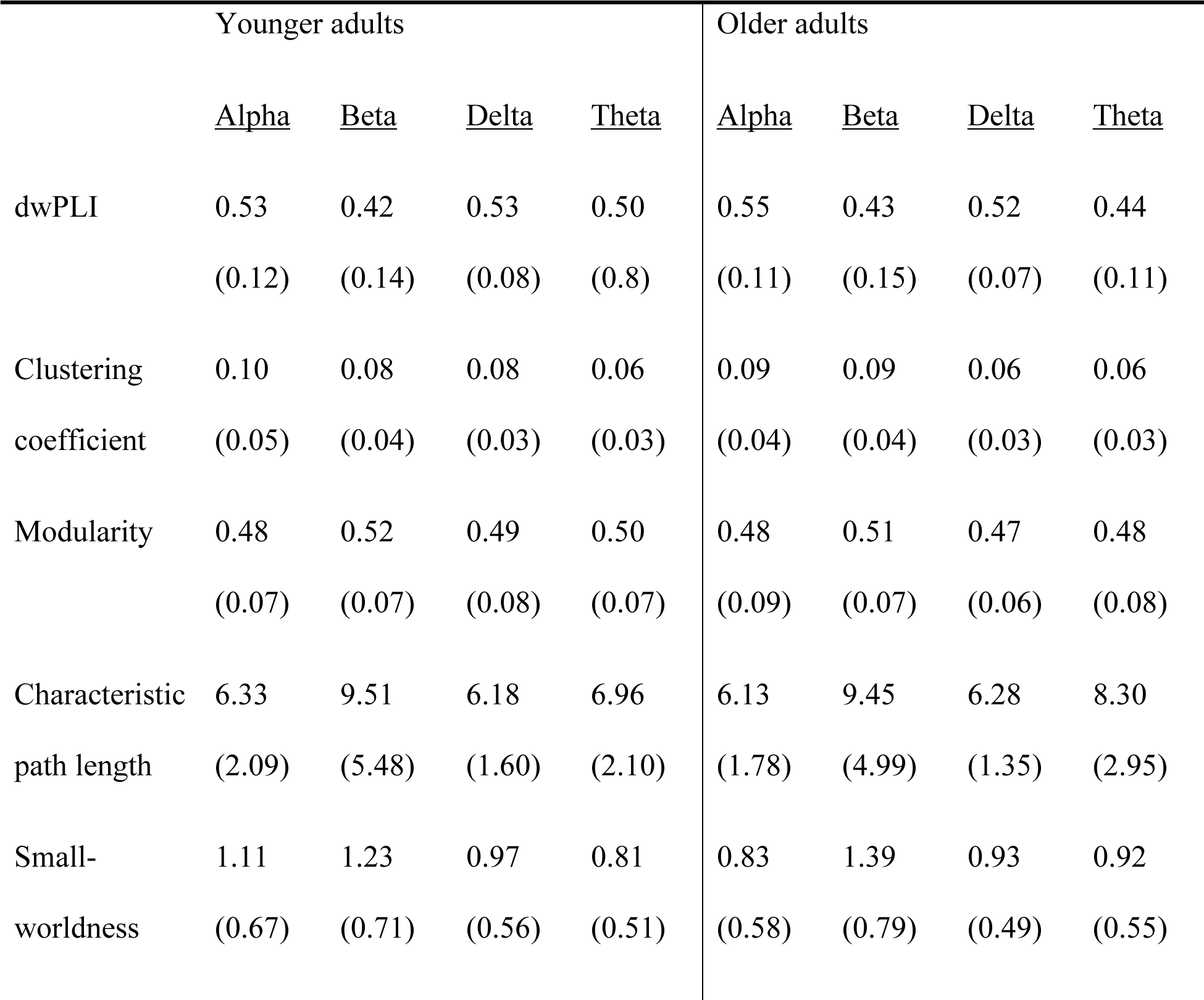
Mean and Standard Deviation for the OMST-Thresholded Graph Theoretical Indices per Age Group.

|  | Younger adults |  |  |  | Older adults |  |  |  |
| --- | --- | --- | --- | --- | --- | --- | --- | --- |
|  | <u>Alpha</u> | <u>Beta</u> | <u>Delta</u> | <u>Theta</u> | <u>Alpha</u> | <u>Beta</u> | <u>Delta</u> | <u>Theta</u> |
| dwPLI | 0.53 | 0.42 | 0.53 | 0.50 | 0.55 | 0.43 | 0.52 | 0.44 |
|  | (0.12) | (0.14) | (0.08) | (0.8) | (0.11) | (0.15) | (0.07) | (0.11) |
| Clustering | 0.10 | 0.08 | 0.08 | 0.06 | 0.09 | 0.09 | 0.06 | 0.06 |
| coefficient | (0.05) | (0.04) | (0.03) | (0.03) | (0.04) | (0.04) | (0.03) | (0.03) |
| Modularity | 0.48 | 0.52 | 0.49 | 0.50 | 0.48 | 0.51 | 0.47 | 0.48 |
|  | (0.07) | (0.07) | (0.08) | (0.07) | (0.09) | (0.07) | (0.06) | (0.08) |
| Characteristic | 6.33 | 9.51 | 6.18 | 6.96 | 6.13 | 9.45 | 6.28 | 8.30 |
| path length | (2.09) | (5.48) | (1.60) | (2.10) | (1.78) | (4.99) | (1.35) | (2.95) |
| Small- | 1.11 | 1.23 | 0.97 | 0.81 | 0.83 | 1.39 | 0.93 | 0.92 |
| worldness | (0.67) | (0.71) | (0.56) | (0.51) | (0.58) | (0.79) | (0.49) | (0.55) |

### Functional Connectivity Strength

First, we hypothesised that age-related decreases in functional connectivity, as measured through dwPLI, would be positively related to age-related word-finding difficulties. Multiple linear regression analysis was used to investigate whether age and dwPLI would predict letter and semantic fluency. The predictors in the delta band explained 13.4% of the variance in semantic fluency scores (*R*^2^ = .10, F(4,95) = 3.66, *p* = .025). Age and the interaction between age and dwPLI were significant predictors of semantic fluency scores *(β* = -2.45, *p* = .020 and *β* = -3.85, *p* = .004, respectively). That is, age-related increases in dwPLI were related to lower semantic fluency scores (see Figure 2). Semantic fluency was not related to an age-related change in dwPLI in any of the other frequency bands, nor was letter fluency related to age-related changes in dwPLI in any of the four frequency bands.

**Figure 2.**
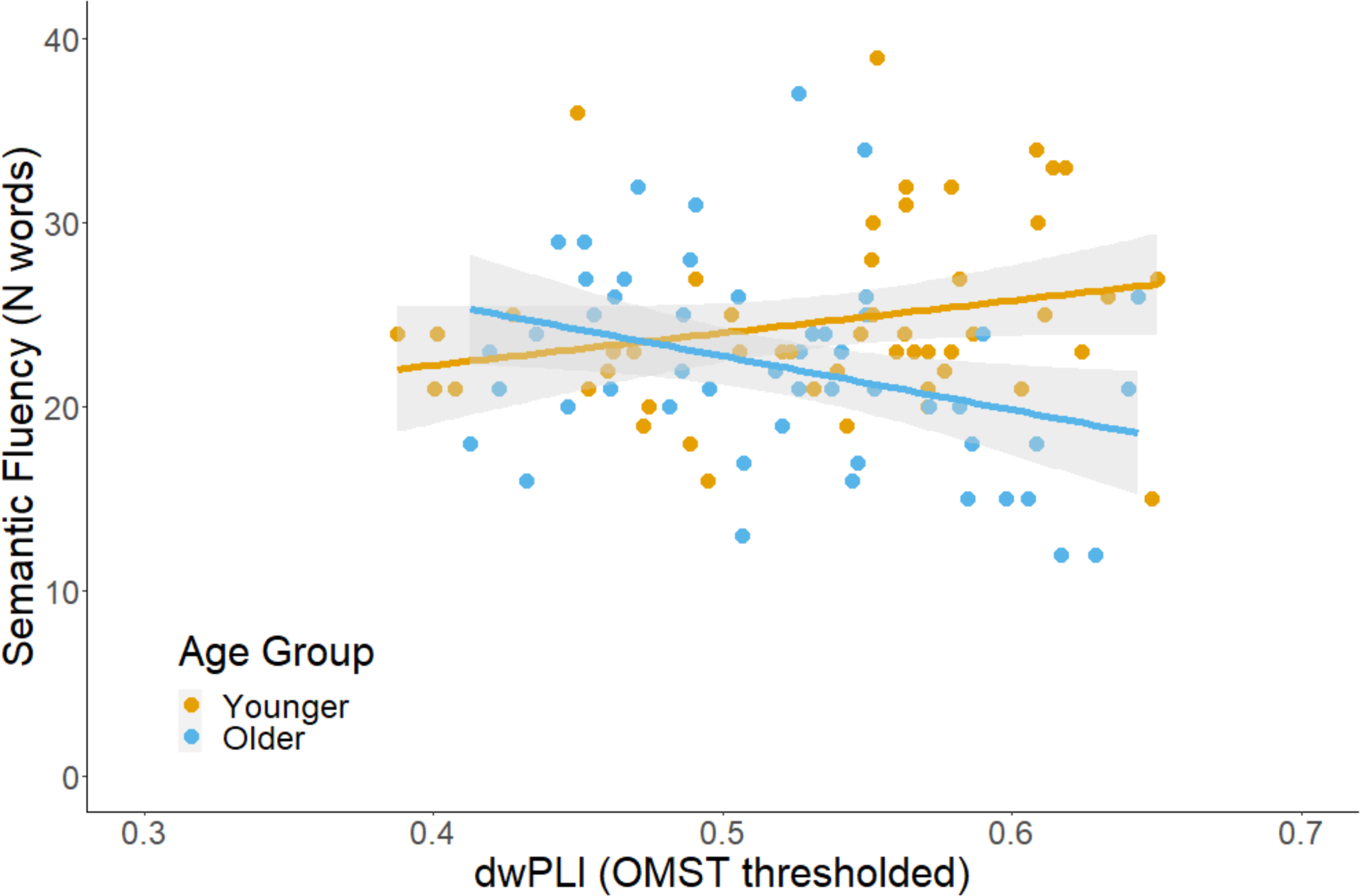
The interaction between age and dwPLI as a significant predictor of semantic fluency in the delta band.

### Brain Segregation

Our second hypothesis was whether age-related decreases in brain segregation, reflecting neural dedifferentiation, are related to reduced word-finding ability. We predicted that age-related changes in brain segregation, as measured through modularity, clustering coefficient, and small world index, would be related to verbal fluency performance. Multiple regression analysis was used to investigate the effect of modularity and clustering coefficient on semantic and letter fluency separately. The predictors in the alpha band explained 14.4% of the variance in semantic fluency scores (*R*^2^ = .08, *F*(6,80) = 2.25, *p* = .047). In the alpha band, both modularity and age independently predicted semantic fluency *(β* = -2.23, *p* = .049 and *β* = -2.83, *p* = .022, respectively). That is, higher modularity scores predicted lower semantic fluency scores, independent of age (see Figure 3). None of the models for letter and semantic fluency indicated that age-related changes in clustering coefficient or modularity predicted letter and semantic fluency scores.

**Figure 3.**
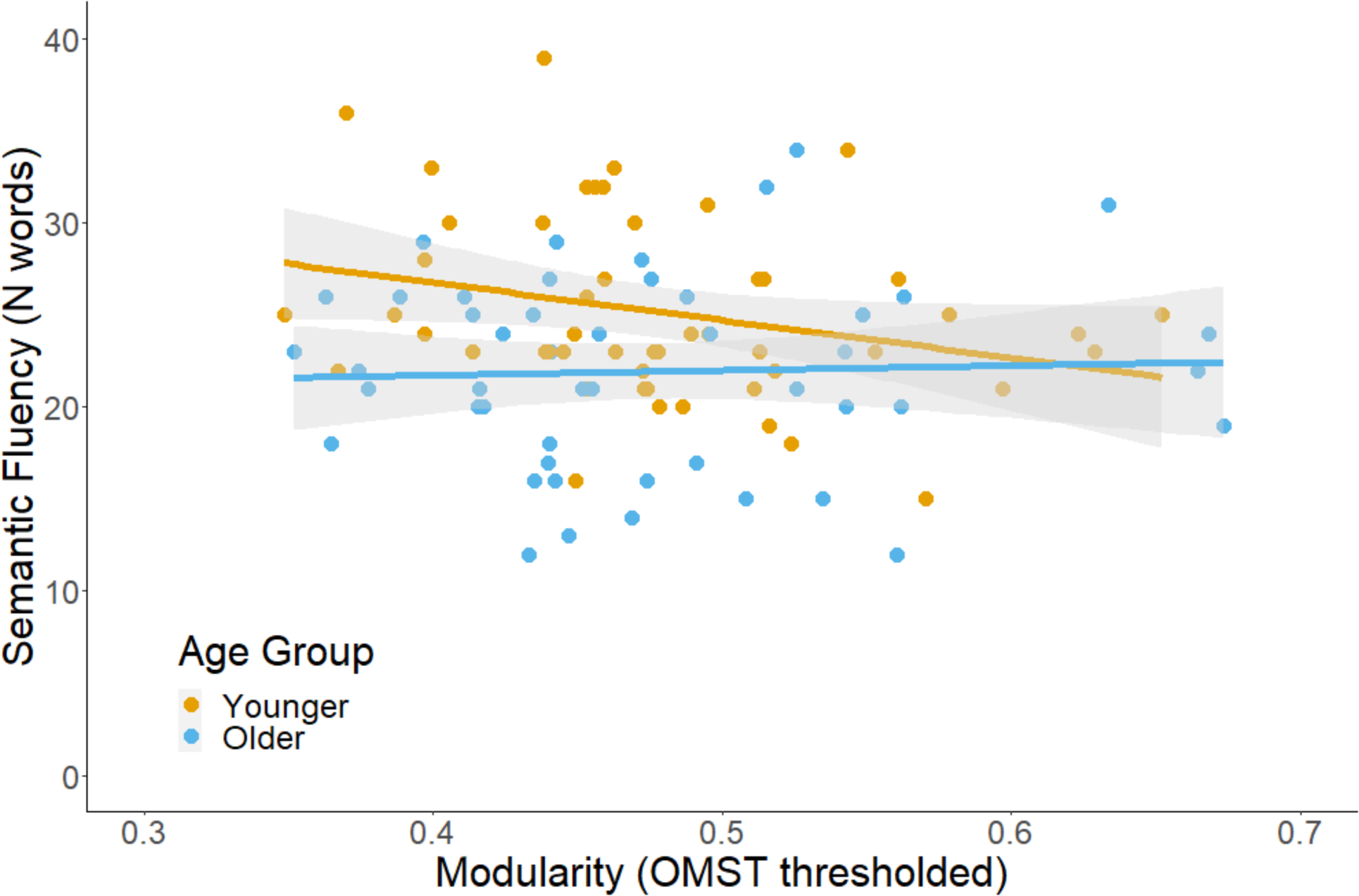
Modularity as a significant predictor, irrespective of age, of semantic fluency in the alpha band.

Because the small-world index suggests a balance between brain integration and segregation, we also ran multiple linear regressions to investigate the effect of age-related changes in the small-world index on verbal fluency performance. In the delta band, the predictors explained 16.6% of the variance in semantic fluency scores (*R*^2^= .12, *F*(4,76) = 3.78, *p* = .007). Both age and the small-world index significantly predicted semantic fluency scores *(β* = -4.08, *p* < .001, and *β* = 2.69, *p* = .041, respectively). That is, a greater small-world index predicted higher semantic fluency scores, irrespective of age (see Figure 4). None of the models for letter and semantic fluency indicated that age-related changes in the small-world index predicted letter and semantic fluency scores.

**Figure 4.**
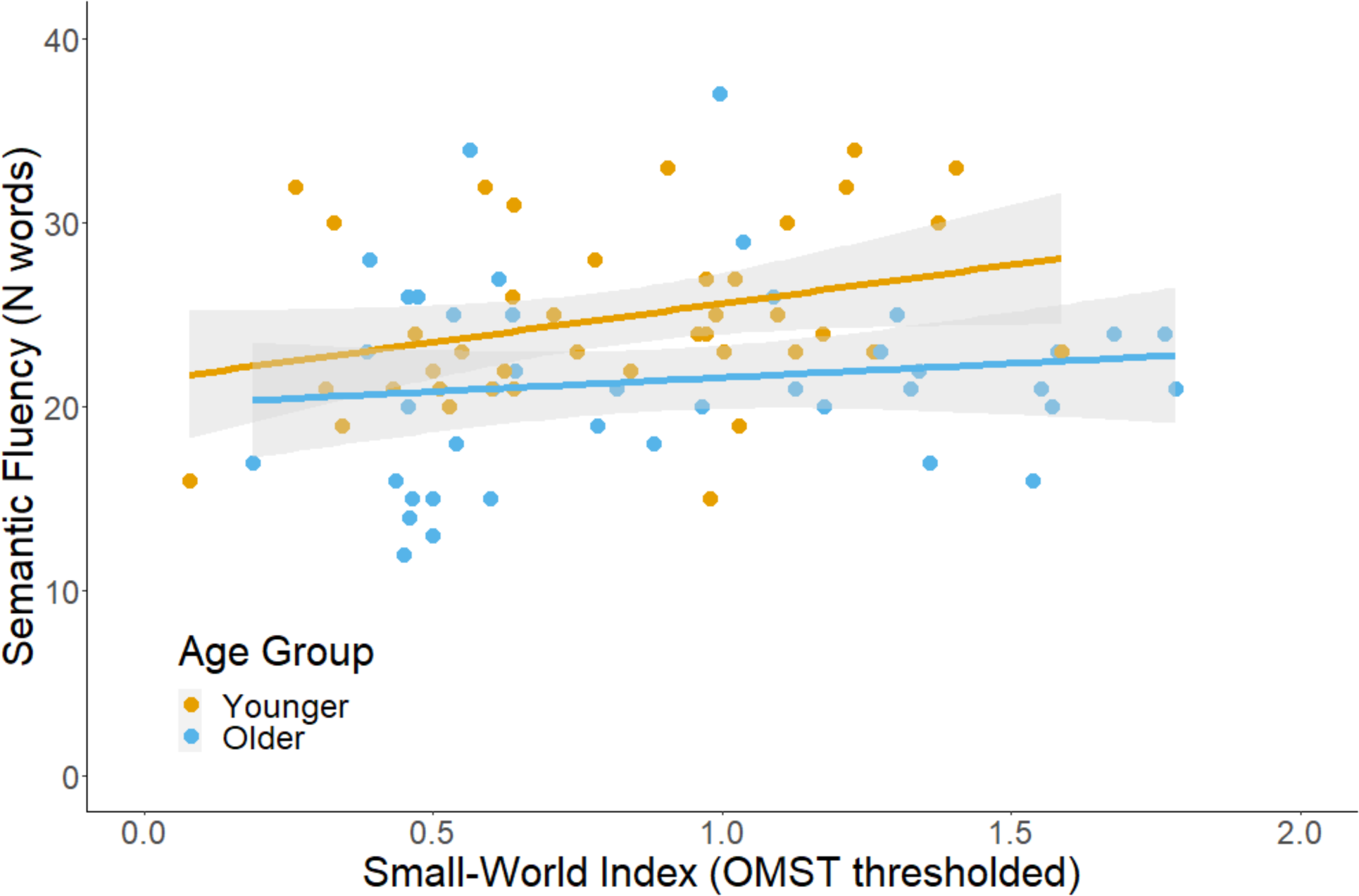
The small-world index as a significant predictor, independently of age, of semantic fluency in the delta band.

### Exploratory Analysis of Brain Integration

Although we did not have a-priori hypotheses about the relationship between age-related changes in brain integration and verbal fluency performance, previous studies linked age-related changes in brain integration measures, such as path length, to changes in cognitive performance (e.g., Stanley et al., 2015). Therefore, we conducted an exploratory analysis (i.e., not preregistered with a-priori hypotheses) using multiple linear regression to investigate the effect of age-related changes in brain integration on verbal fluency performance. In the delta band, the predictors explained 13.2% of the variance in semantic fluency scores (*R*^2^ = .09, *F*(4,94) = 3.57, *p* < .001). Age and the interaction between age and characteristic path length predicted semantic fluency scores *β* = -1.24, *p* = .027 and *β* = 3.64, *p* = .028, respectively). That is, older adults with higher characteristic path length achieved higher semantic fluency scores (see Figure 5). Characteristic path length did not predict semantic fluency in the other frequency bands, nor did it predict letter fluency in any of the frequency bands.

**Figure 5.**
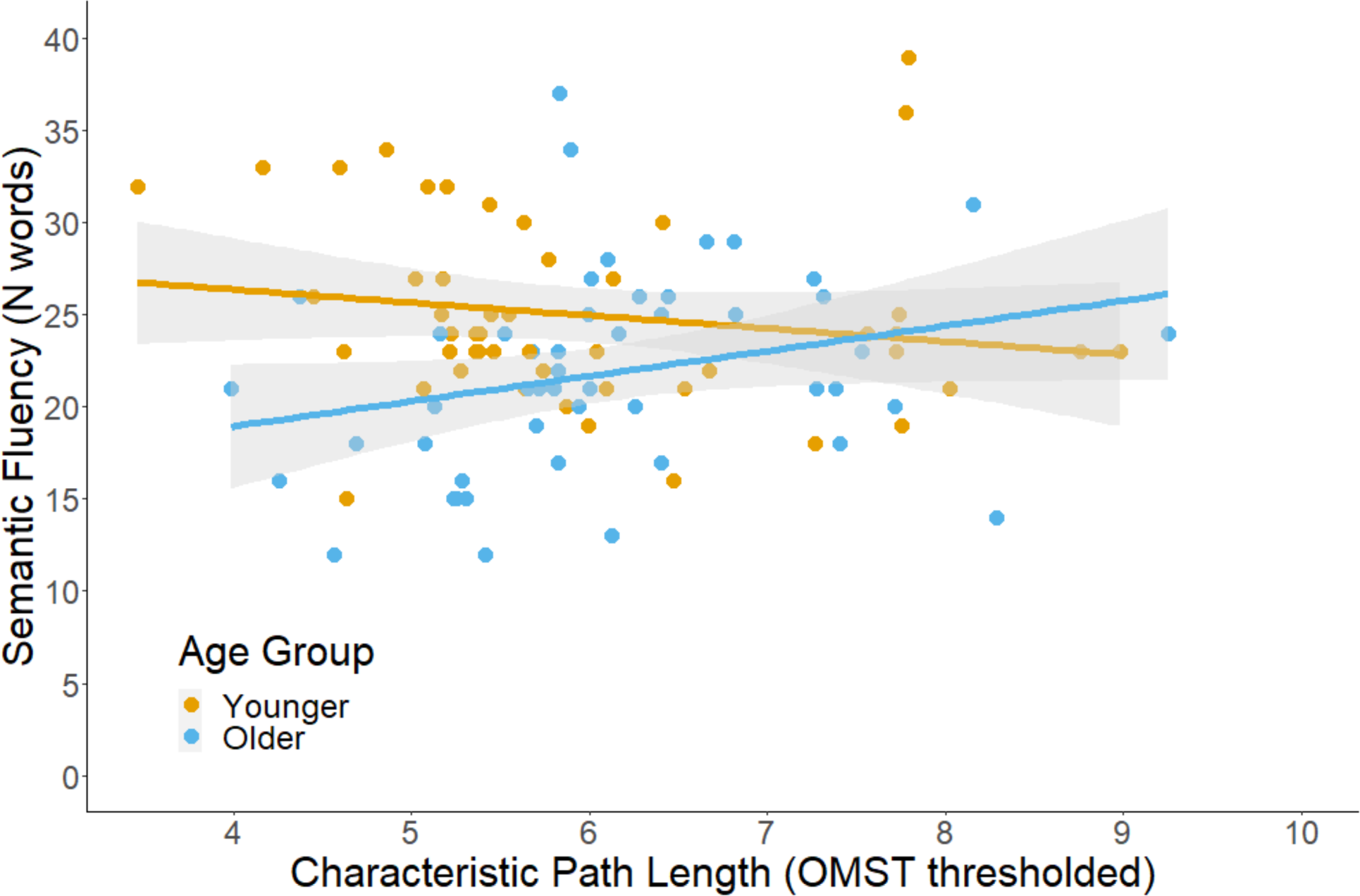
The interaction between age and characteristic path length is a significant predictor of semantic fluency in the delta band.

## Discussion

Using resting-state EEG data, the current study aimed to investigate the relationship between word-finding difficulties in older age and age-related changes in functional brain networks. We hypothesised that age-related decreases in word-finding ability are related to decreases in the connectedness of functional brain networks in older compared to younger adults. We found that, in older adults, greater functional connectedness in the delta band, as measured through dwPLI, was related to lower semantic fluency scores. Several other findings were not in line with our hypotheses. A greater small-world index in the delta band was related to higher semantic fluency performance in both younger and older adults. Greater modularity in the alpha band was related to higher semantic fluency scores, but this was irrespective of age. In an exploratory analysis of brain integration, we found a positive relationship between characteristic path length and semantic fluency in the delta band, only in older adults.

### Age-Related Changes 1n Functional Connectedness and Word-Finding Ability

Our finding that greater functional connectedness related to lower semantic fluency in older adults is in line with previous studies demonstrating age-related changes in the connectedness of functional brain networks (Gaál et al., 2010; Sala-Llonch et al., 2014; Zangrossi et al., 2021), but not with their link with decreases in cognitive performance (Andrews-Hanna et al., 2007; Chow et al., 2022; Fleck et al., 2016). That is, previous studies found that increased connectedness in older adults was related to better cognitive performance. Greater functional connectedness has been hypothesised to indicate greater efficacy of information transfer between different brain areas (Fries, 2005). In this case, increased dwPLI, reflecting overall brain connectedness, would lead to an increase in the brain’s efficiency of information transfer between different brain areas, that is, reduce the slowing of information transfer. Because verbal fluency is a timed task, age-related slowing of information transfer between brain areas in older adults could mean that older adults need more time to access words from their memory. If higher dwPLI values (i.e., greater connectedness) reflect greater efficiency of information transfer between brain areas, one would expect that greater functional connectedness would be related to better word-finding ability. However, we found that greater connectedness in the current study was related to worse word-finding ability.

One possible explanation is that increased dwPLI in older adults represents a pattern of hyper-synchronisation or overload, reflecting noisy communication between different brain regions, and subsequently leading to poorer cognitive performance (Jones et al., 2016; López-Sanz et al., 2017). Our findings for word-finding ability are in line with the studies by Jones et al. (2016) and Lopez-Sanz et al. (2017), who demonstrated that increased resting-state connectivity was related to worse language performance. Moreover, studies in Parkinson’s disease demonstrated that greater functional connectedness, as measured with (dw)PLI in the delta and theta frequency band predicted whether someone had Mild Cognitive Impairment (MCI) or not (Cai et al., 2021; Chaturvedi et al., 2019). Hence, increased delta dwPLI could be explored as a potential marker of pre-onset dementia.

Research has also indicated that individuals with Huntington’s disease exhibit greater delta-band connectivity, which could indicate either pathological or compensatory changes in brain function (Davis et al., 2022). That is, brain activity may be synchronising to a less functional frequency range (here, delta), perhaps as a pathology-related compensatory process, or perhaps reflecting the brain’s inability to inhibit this (potentially less functional) rhythm. Increased delta connectivity may reflect entrainment to a basic resonance property of pyramidal neurons to sustain frequency preference, in contrast to more functional connectivity within task-related oscillatory frequencies (for a discussion on neuronal resonance, see Hutcheon & Yarom, 2000). Resonance plays a crucial role in enabling synchronized activity and oscillatory patterns, and a disruption in the temporal coordination of neuronal activity may lead to cognitive impairments (Lehrer & Eddie, 2013; Uhlhaas & Singer, 2006). In conditions like Alzheimer’s disease, even before disease onset, brain pathology may cause hyperactivity and/or inhibition of neurons, disrupting the neuronal excitation/inhibition (E/I) balance and affecting whole-brain network configurations (Alexandersen et al., 2022; Stam et al., 2023; van Nifterick et al., 2022). Our findings may suggest a similar disruption of the E/I balance in older adults experiencing word-finding difficulties. In this case, neurons exhibit resonance to lower frequencies, specifically the delta range, due to hyper-excitation and/or inhibition of more functional higher frequency ranges, such as beta frequencies.

As an alternative explanation, lower-frequency oscillations, such as delta oscillations, might be the result of compensatory processes to maintain high cognitive performance. With regard to ageing, studies have shown that greater delta band power in younger adults (Mousavi et al., 2020) and greater delta band coherence (i.e., a measure of connectivity) in frontal brain areas in older adults (Fleck et al., 2016) was related to higher semantic fluency scores. In the current study, however, whilst greater delta band connectivity was related to higher semantic fluency scores in younger adults, this was not the case in older adults. In older adults, increased delta connectivity may reflect failed compensatory processes, in line with the idea that the brain is less able to inhibit this lower-frequency oscillatory rhythm. Since our study took a whole-brain approach, we could not determine whether these connectivity effects were region-specific, and other oscillatory patterns may be found when looking at, for example, delta connectivity in frontal brain areas. In younger adults, greater overall connectedness in the delta band might reflect an individual’s ability to inhibit irrelevant responses and maintain internal concentration (Harmony, 2013; Mousavi et al., 2020), which may reverse with age reflecting pathology-related compensatory processes.

### Brain Segregation and Word-Finding Ability Across the Lifespan

Greater modularity in the alpha band predicted lower semantic fluency performance, irrespective of age. Greater brain modularity has been hypothesised to enable greater brain plasticity, because it increases the brain’s efficiency and flexibility to adapt to, for example, age-related anatomical brain changes (Gallen & D’Esposito, 2019). Hence, we would expect that greater modularity would improve word-finding abilities. However, greater modularity was related to lower semantic fluency scores in both younger and older adults, contradicting the suggestion that greater modularity reflects greater brain plasticity and better cognitive functioning. Another explanation is that, to perform verbal fluency tasks, brain integration might be more important than segregation and it is possible that greater brain segregation could be related to lower integration. It has been suggested that greater brain integration is necessary for higher-level cognitive functions, such as language (Bagarinao et al., 2019; Bullmore & Sporns, 2012). However, this seems unlikely as there is a positive relationship between modularity (segregation) and characteristic path length (integration) in the alpha band (see the Supplementary Materials for the analysis). Alternatively, resting-state modularity might perhaps positively relate to some cognitive functions, such as visuospatial working memory, but not others (e.g., numerical working memory; Alavash et al., 2015), or perhaps modularity needs to reach a certain threshold after which it becomes detrimental to cognitive functioning.

Alternatively, one theory proposes that modular brain networks are necessary for quick and simple tasks, whilst complex tasks that require more time benefit more from a lower modular structure (Deem, 2013). For example, greater modularity was negatively related to performance on a complex task, which involved the ability to control attention, whilst modularity was positively related to performance on a simple task (i.e., not involving the control of attention; Yue et al., 2017). Verbal fluency tasks are considered complex tasks as it involves a multitude of cognitive functions to support lexical access (Shao et al., 2014). Hence, in line with the theory by Deem (2013), greater modularity could negatively predict verbal fluency performance. Moreover, our study did not reveal a negative relationship between modularity and letter fluency. The cognitive functions and brain regions underlying letter and semantic fluency are slightly different (Gordon et al., 2018; Shao et al., 2014; Vonk et al., 2019), which could explain the discrepancy between the two tasks in our study.

We also predicted that brain segregation, specifically in the delta band, would play an important role in predicting semantic fluency scores in older adults (Fleck et al., 2016; Mousavi et al., 2020). Fleck and colleagues (2016) argued that maintaining delta band brain segregation in older age is necessary to maintain cognitive performance and decreases will lead to cognitive decline. The current study demonstrated that alpha but not delta brain segregation, as measured through modularity, was related to cognitive performance, and this relationship was irrespective of age. Alpha band activity has been proposed to reflect attention, working memory, and switching abilities (Başar et al., 1999; Stam, 2000). It has been proposed that resting-state alpha band activity is an important indicator of an individual’s readiness for subsequent task performance, potentially through the inhibition of irrelevant pre-task information (Jann et al., 2010; Klimesch et al., 2007). Hence, the current study might indicate that semantic fluency benefits more from a non-modular structure in the alpha band. Less-modular brain networks at rest may allow for a fast reorganisation of the brain network and the direction of attention to the task at hand. Alternatively, the finding in alpha but not delta frequency band modularity might reflect a compensatory mechanism whereby brain activity synchronises within a less functionally specific frequency band (Davis et al., 2022).

### Neural dedifferentiation for word-finding abilities

We hypothesised that brain segregation decreases with age, reflective of neural dedifferentiation. Age-related decreases in brain segregation have been proposed to reflect neural dedifferentiation in older adults (Goh, 2011; Zuo et al., 2017), which means that brain regions and networks become less functionally specific to cognitive processes (Li et al., 2001). Previous studies have supported this idea and showed that greater brain segregation in older adults was related to better memory ability (Chan et al., 2014; Zangrossi et al., 2021). The current study demonstrated an inverse relationship between modularity and word-finding ability, irrespective of age. This is interesting given the expectation that greater modularity would reflect more functional specificity to cognitive functions and, consequently, benefit cognitive functioning. It is possible that modularity does not represent age-related neural dedifferentiation for semantic fluency. Moreover, modularity was computed using resting-state brain networks and not during the semantic fluency task. It is possible that resting-state modularity cannot capture neural dedifferentiation underlying age-related word-finding difficulties.

### The Small-World Index and Word-Finding Abilities

The small-world index is a measure of the organisation of functional brain networks. The current study showed that a greater small-world index in the delta band was related to better semantic fluency performance, irrespective of age. Several studies have linked greater cognitive performance in middle-aged and older adults to a higher small-world index (Douw et al., 2011; Vecchio et al., 2016) and it has been hypothesised that a greater small-world index reflects a more efficient brain (Achard & Bullmore, 2007). A previous study in people with chronic fatigue syndrome demonstrated that the delta band small-world index was negatively related to cognitive dysfunction, which included problems with attention, remembering, and word-finding (Zinn et al., 2017). Our findings add to the literature and indicate that brain networks with small-world topologies could underlie semantic fluency performance in both younger and older adults. Hence, an optimal balance between local and global connectedness might be important for maintaining word-finding ability across the lifespan, irrespective of any age-related decreases.

### Greater Brain Integration is Related to Better Word-Finding in Older Adults

In an exploratory analysis, we investigated the relationship between brain integration and age-related word-finding difficulties. In older adults, a longer characteristic path length in the delta band was associated with higher semantic fluency scores. Shorter characteristic path length would reflect greater global network efficiency because fewer nodes need to be traversed to transfer information from one brain area to another (Bullmore & Sporns, 2009). In contrast, we found that greater characteristic path length related to better semantic fluency in older age. Several studies showed that brain integration decreases with age (Bagarinao et al., 2019; McIntosh et al., 2014; Sullivan et al., 2019), including characteristic path length (Vecchio et al., 2014). Hence, greater characteristic path length (i.e., increased integration) might be necessary to maintain word-finding abilities in older age.

## Limitations and Future Directions

The current study has several limitations. First, the OMST algorithm resulted in too sparse a network in some participants to compute the clustering coefficient and the small-world index. On the one hand, the study was still able to demonstrate the relationship between word-finding and clustering coefficient but, on the other hand, no interaction effects were observed. The latter could be the result of the reduced sample size due to missing values for the clustering coefficient. However, the choice of thresholding is an important one as the incorrect thresholding method can lead to biases and make it difficult to compare across studies. For example, arbitrary thresholding affects the reliability of a study, and bias can appear when one chooses a threshold based on what threshold leads to significant results (Miljevic et al., 2022). In contrast, data-driven thresholding is more objective as the user has no influence on what threshold is chosen. Therefore, we decided to implement the OMST algorithm as it is a data-driven threshold method created to reduce the sparsity of networks, whilst maximising global efficiency (Dimitriadis et al., 2017). Moreover, a recent study indicated that with increasing age, the individual variability in functional brain networks increases (Ma et al., 2021). It is possible that data-driven methods are better suited when comparing functional brain networks, to account for networks that are highly variable between individuals (Bansal et al., 2018). To shed light on functional brain connectivity in ageing, more research is needed using the OMST algorithm for thresholding the brain connectivity graphs so that these studies can be compared.

Another debate in the field of functional brain connectivity obtained through EEG recordings is whether one should project the signals into sensor (i.e., electrodes) or source space (i.e., brain regions underlying the observed electrical activity). Many studies argue for projecting signals into source space as doing so is suggested to resolve problems such as volume conduction and field spread (see Schoffelen & Gross, 2009). However, source-space analyses have their own limitations. For example, there are multiple methods for identifying the underlying sources and the estimation of parameters needed for source localisation is very complex and relies on a number of assumptions (Mahjoory et al., 2017; Miljevic et al., 2022). In addition, a recent study showed that sensor space might be more suitable for conducting functional brain connectivity analyses as brain network indices, such as the characteristic path length, can change after projecting brain activity into source space (Koutlis et al., 2021).

Moreover, the current study aimed to conduct whole-brain network analyses and we did not have a-priori hypotheses about the underlying brain regions. Whole-brain network analyses are important to understand the effect of age on how functional networks combine the information processed by the brain (Geerligs et al., 2015). To reduce the influence of volume conduction and field spread. The debiased weighted Phase Lag Index was used to compute the connectivity between the EEG sensors (Lai et al., 2018; Vinck et al., 2011). Hence, the issues created by volume conduction were addressed without the need to conduct source localisation in the current study. Nevertheless, future studies could build on the results of the current study by investigating the sources underlying the functional brain connectivity patterns we have observed.

Finally, it is important to note that the current study identified brain networks from resting-state data, hence, these networks were not obtained during the verbal fluency tasks. Functional brain networks underlying verbal fluency tasks may yield different patterns from resting-state data, and age-related changes might be reflected differently in task-dependent functional connectivity. However, resting-state EEG analyses have been argued to be informative (Rosazza & Minati, 2011; van Diessen et al., 2015) and such analyses can be useful in providing insights into cognition, and the diagnosis, development, and treatment of neurodegenerative diseases (Ishii et al., 2017; O’Neill et al., 2018).

The current study also identified some gaps and recommendations for future research. First, we demonstrated that increased functional connectedness in the delta band was related to age-related word-finding difficulties. Future studies should explore this relationship among healthy older adults, adults at risk of dementia, and adults in the beginning stages of dementia. Investigate whether such a measure, in conjunction with neuropsychological assessments, could contribute to the early detection of cognitive impairment and dementia. Second, interventions could be developed that aim to increase delta band functional connectedness as this could increase neuroplasticity and, consequently, improve cognitive outcomes in older adults. Third, considering our contrasting finding that increased modularity was related to poorer word-finding ability irrespective of age, future studies could investigate when the extent of brain segregation becomes detrimental to word-finding ability. Additionally, examining potential differences in this effect between younger and older adults would be valuable. Finally, because modularity has been proposed to be important in predicting neuroplasticity outcomes following intervention, such as cognitive training (Gallen & D’Esposito, 2019), it is important to investigate the interactive effect between modularity, complexity of cognitive functioning, and intervention outcomes. Such research is essential not only in ageing populations but also in those with neurodegenerative diseases.

## Conclusion

The current study investigated the link between functional brain connectivity and word-finding abilities in younger and older adults. We found that changes in functional brain connectivity, such as in overall connectedness and characteristic path length, related to worse performance on semantic fluency, but only in older adults. Modularity and the small-world index also predicted semantic fluency performance, but this was irrespective of age. Moreover, changes in functional brain connectivity were specific to the frequency band, possibly reflecting changes in cognitive control and the ability to inhibit irrelevant responses or a compensatory shift to less functionally specific frequency bands. This is the first study demonstrating that age-related word-finding difficulties can be linked to changes in functional brain connectivity.

## Author Note

The study hypotheses, design, and statistical analyses were preregistered on the Open Science Framework (https://osf.io/u6p42). MATLAB and R code, Cognitive Reserve questionnaires, the fully anonymised dataset, and additional online materials will be made openly available after publication.

Authors report no conflict of interest.

## Funding sources

This work was supported by a Faculty of Science & Technology scholarship from Lancaster University awarded to Elise J. Oosterhuis, and a Biotechnology and Biological Science Research Council (grant number BB/S008527/1) awarded to Helen E. Nuttall which supported Kate Slate.

## Notes

https://osf.io/u6p42

### Competing Interest Statement

The authors have declared no competing interest.

